# *Anapc5* and *Anapc7* as genetic modifiers of KIF18A function in fertility and mitotic progression

**DOI:** 10.1101/2024.12.03.626395

**Authors:** Carleigh Nesbit, Whitney Martin, Anne Czechanski, Candice Byers, Narayanan Raghupathy, Ardian Ferraj, Jason Stumpff, Laura Reinholdt

**Affiliations:** The Jackson Laboratory, Bar Harbor, ME 04609; Department of Molecular Physiology and Biophysics, University of Vermont, Burlington, VT; The Roux Institute at Northeastern University, Portland, ME

## Abstract

The kinesin family member 18A (*KIF18A*) is an essential regulator of microtubule dynamics and chromosome alignment during mitosis. Functional dependency on KIF18A varies by cell type and genetic context but the heritable factors that influence this dependency remain unknown. To address this, we took advantage of the variable penetrance observed in different mouse strain backgrounds to screen for loci that modulate germ cell depletion in the absence of KIF18A. We found a significant association at a Chr5 locus where anaphase promoting complex subunits 5 (*Anapc5*) and 7 (*Anapc7*) were the top candidate genes. We found that both genes were differentially expressed in a sensitive strain background when compared to resistant strain background at key timepoints in gonadal development. We also identified a novel retroviral insertion in *Anapc7* that may in part explain the observed expression differences. In cell line models, we found that depletion of KIF18A induced mitotic arrest, which was partially rescued by co-depletion of ANAPC7 (APC7) and exacerbated by co-depletion of ANAPC5 (APC5). These findings suggest that differential expression and activity of *Anapc5* and *Anapc7* may influence sensitivity to KIF18A depletion in germ cells and CIN cells, with potential implications for optimizing antineoplastic therapies.

## INTRODUCTION

Cell cycle progression is governed by highly conserved proteins and pathways that monitor and control DNA repair/replication, chromosome alignment, separation of sister chromatids, and accurate chromosome segregation. During M-phase, the spindle assembly checkpoint (SAC) ensures the fidelity of chromosome segregation by preventing the E3 ubiquitin ligase, anaphase promoting complex (APC/C) from degrading the proteins that affix sister chromatids until all of the replicated chromosomes are properly attached to spindle microtubules (McAinsh and Kops 2023). However, some chromosome attachment defects, including merotelic attachments, can escape detection, leading to aneuploidy and chromosome instability (CIN) (Kops, Weaver et al. 2005).

The kinesin family member 18A (*Kif18a*) is one of a number of kinesin proteins that control kinetochore microtubule dynamics and regulate chromosome alignment during mitosis and meiosis (Mayr, Hummer et al. 2007, Stumpff, von Dassow et al. 2008). *Kif18a* is also one of a few cell cycle genes that are dispensable for development but are uniquely required for cell cycle progression in germ cells (Whitney, Royle et al. 1996, Nadler and Braun 2000, Agoulnik, Lu et al. 2002, Meetei, Sechi et al. 2003, Liu, Zhao et al. 2010, Hartford, Luo et al. 2011). Loss of function mutations in *Kif18a* cause M-phase arrest in mitotically and meiotically dividing germ cells (Ward, Reinholdt et al. 2003, Reinholdt, Munroe et al. 2006, Mihajlovic, Byers et al. 2023). In human cells, KIF18A is also critical for proliferation of neoplastic cells that exhibit CIN, including HeLa cells (Marquis, Fonseca et al. 2021). Like germ cells, HeLa cells also exhibit mitotic arrest in the absence of KIF18A. However, somatic cells and diploid cell lines like RPE1 cells with similar loss of KIF18A progress through mitosis despite apparent chromosome instability (Czechanski, Kim et al. 2015).

Cellular context is a key aspect of mitotic progression in cells lacking KIF18A, however genetic context is also important. Loss of KIF18A function in mice leads to recessive infertility with variable penetrance, ranging from 90% penetrance on the wild derived inbred strain background, CAST/EiJ (CAST) to 37% penetrance on the common laboratory inbred strain background C57BL/6J (B6) (Reinholdt, Munroe et al. 2006). These data provide evidence that there are genetic factors that limit or promote cell cycle progression in germ cells in the absence of functional KIF18A. Discovery of these factors would explain key aspects of genomic surveillance in the germ line and may also reveal molecular pathways that allow for cell cycle progression of cancer cells with CIN.

To discover the genetic factors regulating cell cycle progression in germ cells lacking KIF18A function, we used genetic association mapping to statistically link variation in germ cell populations to potential modifier loci in the genomes of F2 mice generated from B6 and CAST crosses. We identified a significant association to a locus on Chr5 containing two components of the APC/C, *Anapc7* and *Anapc5*. We found that expression levels of these genes during the proliferative stages of germ cell development differ significantly depending on the gonadal sex and strain (CAST vs. B6) suggesting that, in part, the underlying genetic variation found in these genes has downstream consequences on gene expression, which may in turn influence the functional availability of APC/C. Consistent with this model, there are genetic variants in both genes that could impact both the expression level and the function of ANAPC7 (APC7) and ANAPC5 (APC5) proteins, including a germ line-expressed retroviral element just upstream of the *Apc7* promoter that is unique to the CAST background. We also found that cell cycle arrest induced by depletion of KIF18A in a cell model could be alleviated by co-depletion of APC7 and exacerbated by co-depletion of APC5, providing further support that natural genetic variation influencing transcript abundance in these two genes regulates cell cycle progression in the context of chromosome alignment defects.

## RESULTS

Laboratory mice carrying a loss of function mutation (*gcd2*, R308K) in *Kif18a* are infertile due to mitotic arrest of developing germ cells (Reinholdt, Munroe et al. 2006, Liu, Zhao et al. 2010, Czechanski, Kim et al. 2015). We previously showed that infertility was 67% penetrant in *Kif18a* deficient (*Kif18a^gcd2/gcd2^*) F2 progeny from an intersubspecific cross between the classic inbred strain C57BL/6J (B6) and the wild-derived inbred strain CAST/EiJ (CAST). Subsequent backcrossing of F2 mice to the CAST background increased the penetrance of infertility to 90%, while subsequent backcrossing to B6 decreased the penetrance to just over 37%. In all cases, fertility status was directly related to the degree of germ cell depletion, which was quantified by direct counting of labeled germ cells in whole mount embryos, fetal gonads, histological sections of ovaries (ovarian follicles), or by estimating the percentage of seminiferous tubules exhibiting germ cell loss in juvenile and adult testis (Reinholdt, Munroe et al. 2006). In *Kif18a* mutant mice, germ cell depletion is apparent from the time germ cells populate the fetal, bipotential gonad through to sexual maturity and is concurrent with mitotic arrest of gonial cells (spermatgonia and oogonia). Additional roles for *Kif18a* during meiosis in the oocyte have also been demonstrated (Tang, Pan et al. 2018, Mihajlovic, Byers et al. 2023). The variable penetrance of these phenotypes suggests that in dividing germ cells, functional dependency on *Kif18a* is regulated by unknown genetic modifiers.

We sought to map these genetic modifiers using the germ cell population as a quantitative phenotype. To generate a large cohort of F1 animals, we used *in vitro* fertilization with cryopreserved sperm from congenic CAST.129-*Kif18a^gcd2/+^* males and oocytes from B6 females. The resulting heterozygous F1 progeny were intercrossed to produce 2,281 F2 progeny. From these, gonads were collected for immunohistochemistry from 477 B6;CAST-*Kif18a^gcd2/gcd2^* (*gcd2/gcd2*) and 93 B6;CAST-*Kif18a^+/+^* (+/+) controls at P18 (18 days post-partum) (Figure 1A). Consistent with our previous results, the germ cell population in *gcd2/gcd2* F2 gonads was significantly depleted compared to controls, but the severity of germ cell depletion showed interindividual variability (Figure 1B). Germ cell populations in the testes were more severely affected (Figure 1B), but this difference was at least partly driven by the technical limitations of histopathological quantification of germ cells in testis.

**Figure 1.**
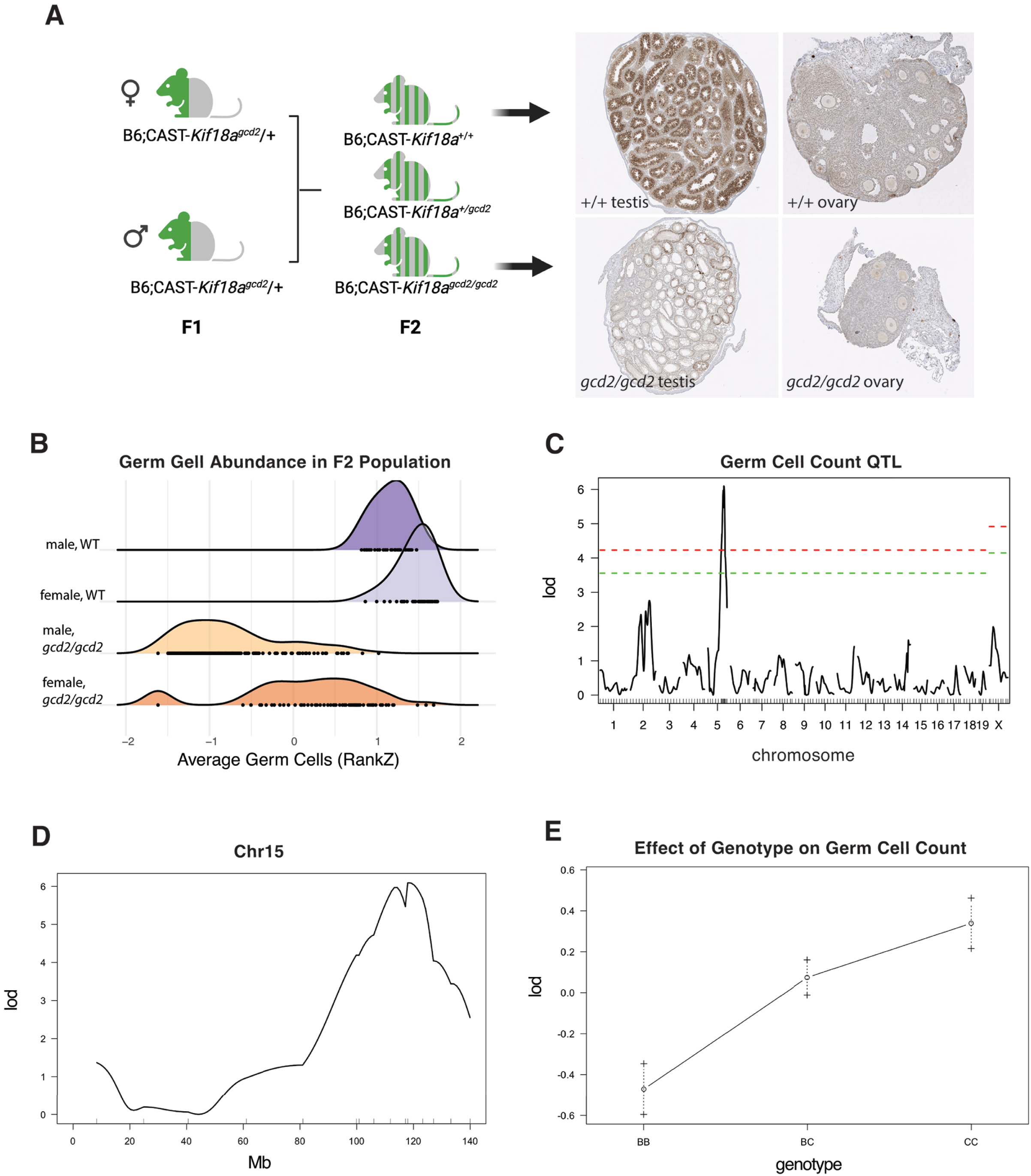
Genetic mapping of a *Kif18a* modifier locus in laboratory mice reveals a locus on Chr5. F2 mapping cross and examples of histological sections with labeled germ cells (A). RankZ transformed germ cell population (sampled from 4 step sections per individual B6;CAST- *Kif18a^gcd2/gcd2^* (n=477) and B6;CAST-*Kif18a^+/+^*(N=93) F2 progeny (B). Genome-wide association map of average germ cell population (inverse normal transformation) in B6;CAST-*Kif18a^gcd2/gcd2^*F2 progeny with sex as a covariate, red=0.01 and green=0.05 significance thresholds based on 1000 permutations (C). LOD scores across Chr 5 locus (D). Transgressive allele effects at rs3670250 (E).

We performed genome wide association mapping of germ cell quantity in a subset (225) of the *gcd2/gcd2* F2s and found a quantitative trait locus (QTL) on Chr5 with a LOD (logarithm of the odds) score exceeding the 0.05 and 0.01 significance thresholds (Figure 1C). After genotyping with additional SNP markers to narrow the interval, we found that the SNP marker (rs3670250) with the peak LOD score (6.09) was at chr5:119,982,112 (GRCm38) (Figure 1D). The allele effects at the peak SNP were additive and opposing with respect to the parental phenotypes (i.e. transgressive) (Figure 1E). To evaluate candidate genes in the interval, we generated transcriptome data from B6 and CAST testis (P5) and fetal ovaries (11.5-13.5 dpc), focusing on developmental time points where mitotically dividing germ cells are enriched in the gonad. We also took advantage of publicly available transcriptome data sets from adult B6 and CAST testes. Using these data, we narrowed the gene list to only those genes expressed in testes and ovaries, with the rationale that the best candidate modifier would be a gene expressed in both tissues. Since B6 is the mouse reference genome and short-read genome assemblies of CAST are published, we used publicly available, reference-based variant (SNP/INDEL) call sets and narrowed the candidate gene list to genes with protein damaging variants, and further to those genes with functional annotations (GO) and/or phenotype annotations that are relevant to the known functions and phenotypes associated with *Kif18a* depletion. This resulted in a short list of candidate genes (*Anapc5, Anapc7, Cit, Kntc1, Pebp1, Rbm19, Setd1b, Tbx3, Trpv4*) (Supplementary Table 2). Because every gene in the interval also has non-coding variation that could plausibly influence differential gene expression in B6 and CAST, we also looked for genes that showed significant differential expression between B6 and CAST in either testes (P5 and adult) or fetal ovary (Supplementary Table 3) at developmental timepoints when mitotically dividing germ cells are enriched. We found that all nine showed significant differential expression in at least one of the tissues / timepoints that we examined.

We focused on two candidate genes, *Anapc5* (*Apc5*) and *Anapc7* (*Apc7*), given their functional role in the anaphase-promoting complex (APC), which is central to mitotic progression and kept inactive by the spindle assembly checkpoint (SAC) until all kinetochores are attached to mitotic spindle microtubules. These two candidates were also of interest because the dependency of certain cancer cell lines on *Kif18a* is affected by APC activity (Gliech, Yeow et al. 2024). Gleich et al. demonstrated that cancer cell lines are uniquely sensitive to pharmacological KIF18A inhibition due to weakened spindle assembly checkpoints, are intolerant to reduction in APC/C activity, and exhibit co-dependency on certain APC/C subunits (APC1, APC4, and APC8). These studies support a model where the balance between SAC signal and APC/C strength drives sensitivity to KIF18A deficiency and highlight the importance of APC/C member stoichiometry to relative APC/C strength (Gliech, Yeow et al. 2024).

We interrogated publicly available variant data for laboratory mouse strains (Ball, Bogue et al. 2024) and found that CAST *Apc5* and *Apc7* alleles harbor protein-coding variants. Additionally, the CAST *Apc5* allele contains a potentially deleterious, predicted stop-gain variant (rs252980450) that impacts a minor transcript (ENSMUST00000199406.5). Our gene expression data revealed significantly increased expression of *Apc7* in CAST testes (P5 and adult), but not fetal ovary (Figure 2A, Supplementary Figure 1). We found that *Apc5* expression was significantly lower in CAST fetal ovaries, but not in testes (Figure 2A, Supplementary Figure 1). These data suggest that differences in the stoichiometry of these two APC subunits could influence mitotic progression of KIF18A deficient germ cells, although the underlying mechanism may differ depending on the developmental program (male vs. female).

**Figure 2.**
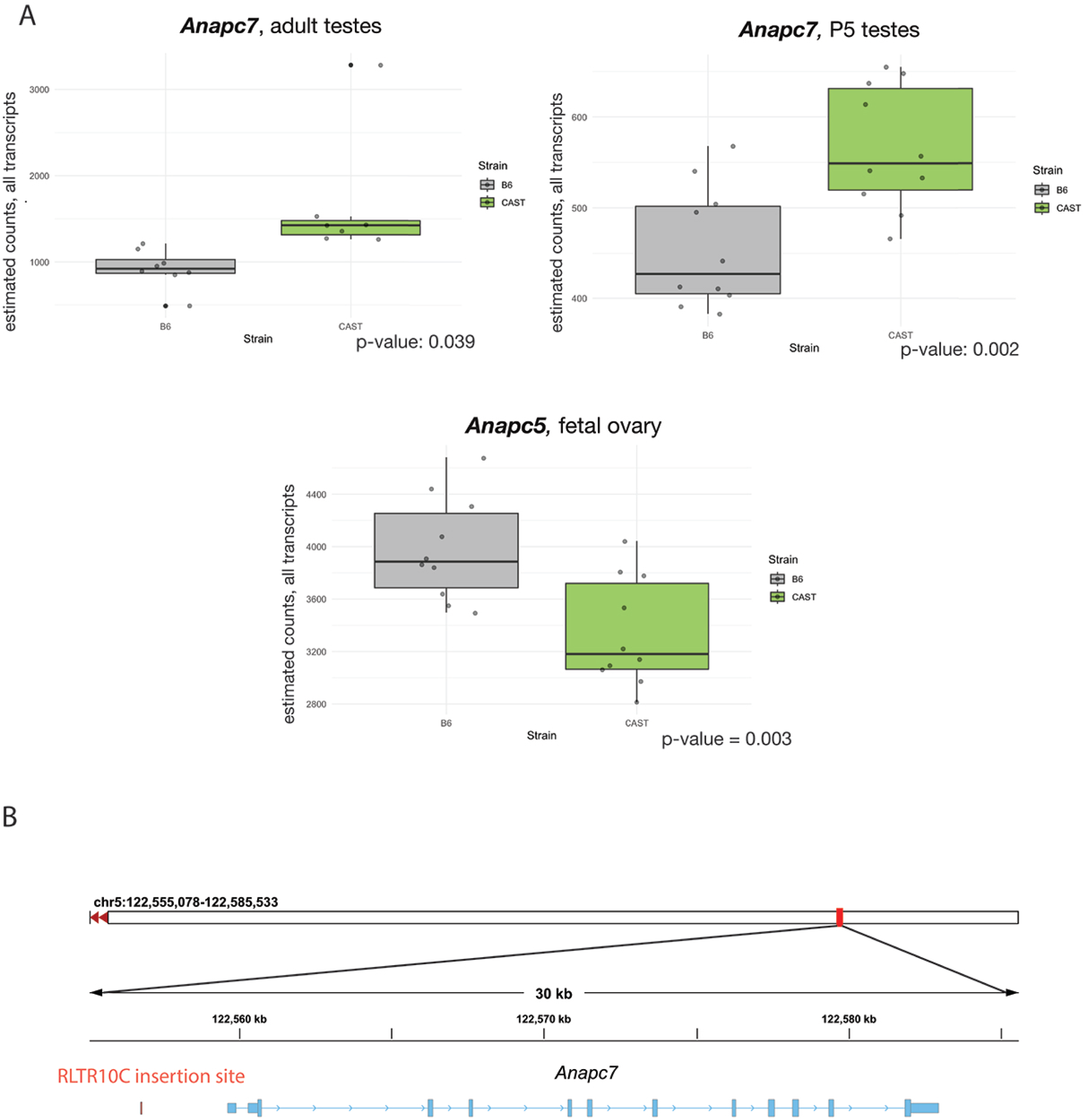
Differential expression and allelic variation in CAST and B6 mice. *Anapc7* (Apc7) and *Anapc5 (Apc5)* expression level data from C57BL/6J and CAST/EiJ juvenile testes (5 dpp), adult testes, and fetal ovary (n=5 biological replicates per tissue type/strain) (A). Genome browser view of *Anapc7 (Apc7)* with upstream site of endogenous retroviral RLTR10C insertion (B).

Because current mouse variant databases are primarily populated by SNPs and INDELs from short-read assemblies, we took advantage of recently published variant call sets derived from long-read genome assemblies to further explore structural variants that might underlie these gene expression differences. While we did not find structural variants in the coding regions of either of these genes, we did identify a previously uncharacterized endogenous retroelement insertion upstream of the *Apc7* start site in CAST. This ∼4 kb insertion is an endogenous retrovirus (ERV), RLTR10C (Figure 2B), a subtype of retroelements that act as transcriptional enhancers in the male germ line (Sakashita, Maezawa et al. 2020). The presence of this element may explain the increased expression of *Apc7* observed in CAST/EiJ testes.

To determine how altered levels of APC5 and APC7 affect the sensitivity of cells to loss of KIF18A activity, the impact of co-depleting APC5 or APC7 with KIF18A was assessed in multiple immortalized cancer cell lines known to have varying sensitivity to depletion of KIF18A (Marquis, Fonseca et al. 2021, Payton, Belmontes et al. 2024). Knockdown of all proteins of interest (KIF18A, APC5, and APC7) was confirmed by Western Blot (Figure 3 & 4, Supplementary Figure 2). All CIN cell lines evaluated (HeLa and two ovarian cancer cell lines; SKOV3 and OVCAR3) demonstrated a significant mitotic arrest with depletion of KIF18A (p <0.0001 for all cell lines). The same effect was not seen in retinal pigmented epithelial cells (RPE1), a near diploid cell line (Figures 3 & 4). In RPE1, HeLa, and OVCAR3 cell lines, there was no appreciable difference in mitotic index between controls and cells depleted of APC7. In SKOV3 cells, the mitotic index of cells depleted of APC7 was lower than that of controls (1.8% vs 4.8%, p <0.0001). In the ovarian cell lines sensitive to KIF18A depletion (SKOV3, and OVCAR3), a partial rescue of the mitotic arrest phenotype was observed when APC7 was co-depleted (SKOV3, p=0.036 and OVCAR3, p=0.0025), however a significant change in the mitotic arrest caused by KIF18A depletion was not observed in HeLa or RPE1 cells co-depleted of APC7 (Figure 3). In RPE1 cells and OVCAR3 cells, there was no appreciable difference in mitotic index between controls and cells depleted of APC5, however, a mild increase in mitotic index was seen in SKOV3 cells (4.9% vs 3.0%, p=0.008) and HeLa cells (8.2% vs 5.3%, p<0.0001) depleted of APC5 compared to controls. An exacerbation of the mitotic arrest phenotype was seen following co-depletion of KIF18A and APC5 in HeLa cells (p < 0.0001)(Figure 4). Taken together, these data indicate that the mitotic arrest caused by KIF18A inhibition can be enhanced by loss of APC5 and reduced by loss of APC7 in some cell types.

**Figure 3:**
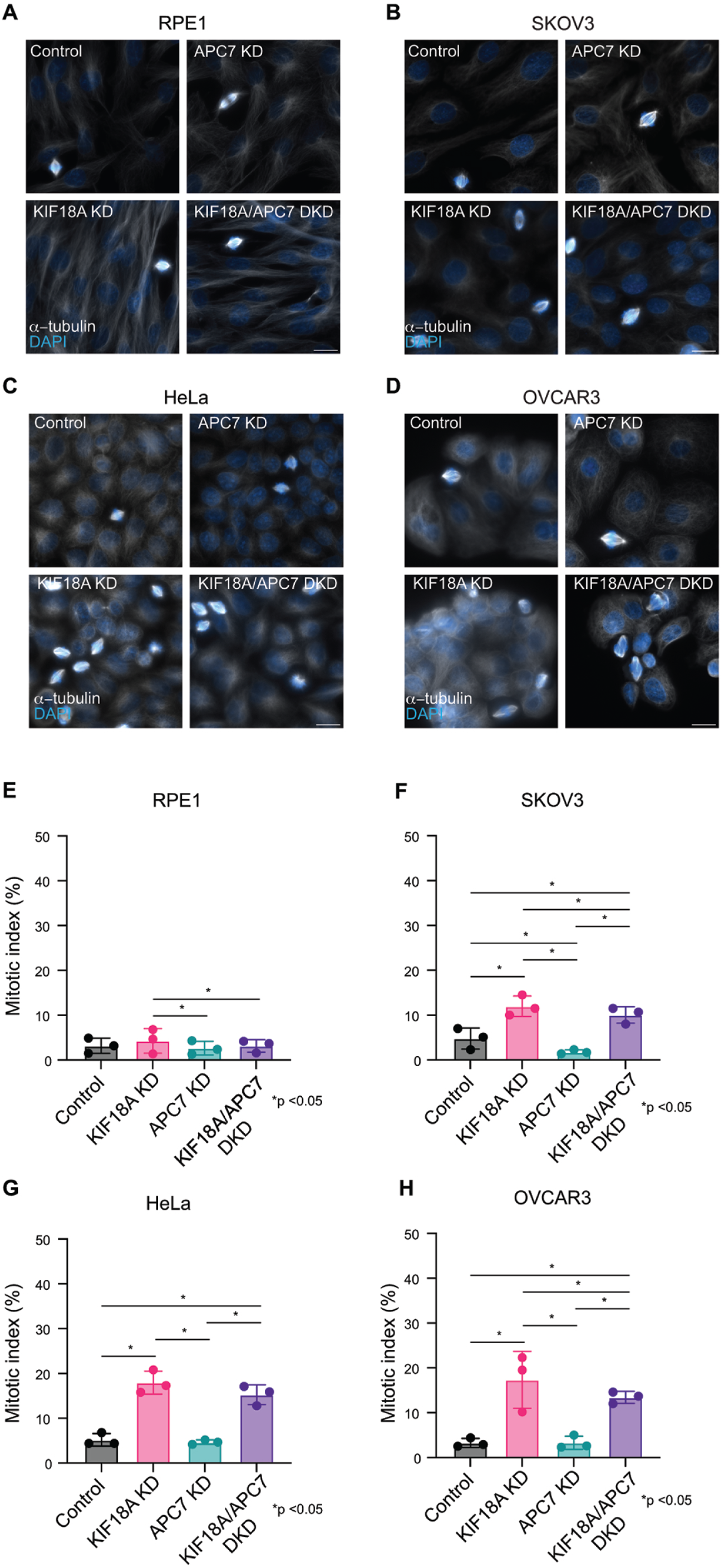
Co-depletion of APC7 with KIF18A leads to partial rescue of mitotic arrest phenotype in human cells. A,B,C,D: Representative images from RPE1 (A), SKOV3 (B), HeLa (C) and OVCAR3 cells (D) treated with control, *KIF18A*, and/or *APC7* siRNAs. DNA (DAPI, blue) and microtubules (α-tubulin, white) are labeled. Scale bars are 20 microns. E,F,G,H: Percentage of mitotic cells (mitotic index) observed in fixed populations of RPE1 (E), SKOV3 (F), HeLa (G), and OVCAR3 (H) cells treated with control, *KIF18A*, and/or *APC7* siRNA. n = 3 independent biological samples per condition for each cell type. For HeLa n=12,399 cells analyzed, for RPE1, n=5,844 cells analyzed, for SKOV3 n=5,344 cells analyzed, for OVCAR3 n=5718 cells analyzed. Statistical analysis performed using chi squared tests.

**Figure 4:**
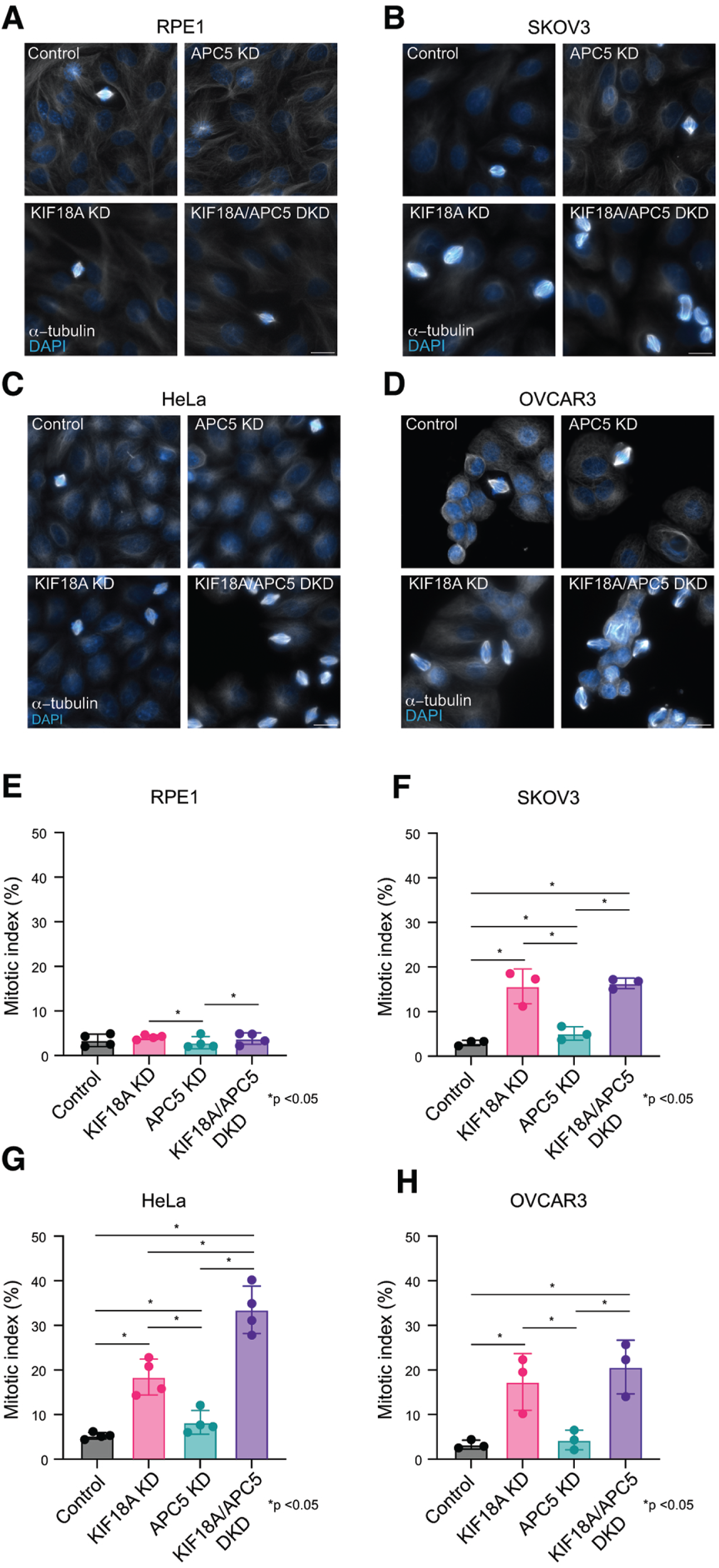
Co-depletion of APC5 with KIF18A leads to exaggeration of mitotic arrest phenotype. A,B,C,D: Representative images from RPE1 (A), SKOV3 (B), HeLa (C) and OVCAR3 cells (D) treated with control, *KIF18A*, and/or *APC5* siRNAs. DNA (DAPI, blue) and microtubules (α-tubulin, white) are labeled. Scale bars are 20 microns. E,F,G,H: Percentage of mitotic cells (mitotic index) observed in fixed populations of RPE1 (E), SKOV3 (F), HeLa (G), and OVCAR3 (H) cells treated with control, *KIF18A*, and/or *APC5* siRNA. n = 4 independent biological samples per condition for HeLa and RPE1; n = 3 for SKOV3 and OVCAR3. For HeLa n=16,335 cells analyzed, for RPE1, n=11,493 cells analyzed, for SKOV3 n=5,365 cells analyzed, for OVCAR3 n=5,608 cells analyzed. Statistical analysis performed using chi squared tests.

## DISCUSSION

Loss of function mutations in *Kif18a* cause mitotic chromosome misalignment and kinetochore-microtubule attachment defects, triggering M-phase arrest in germ cells and in chromosomally unstable cell lines. In laboratory mice, these cell cycle defects lead to infertility with variable expressivity and penetrance depending on sex and genetic background (Ward, Reinholdt et al. 2003, Reinholdt, Munroe et al. 2006). Here, we sought to exploit these laboratory mouse strain background differences to identify the genetic variants that allow cells to escape these cell cycle defects. Through quantitative trait mapping, we identified two subunits of the anaphase promoting complex (APC) as potential candidate regulators (*Apc5* and *Apc7*). The APC is an E3 ubiquitin ligase that is active through much of the cell cycle and promotes progression through mitosis by degrading regulators at specific time points, particularly at the metaphase to anaphase transition. In vertebrates, the APC consists of 14 different subunits which are organized into three domains; the “platform,” the “catalytic core,” and the “tetratricopeptide repeat” or TPR lobe. *Apc7* is part of the TPR lobe which comprises the majority of the mass of the APC and acts as a scaffold as well as mediator for interactions with regulatory proteins. *Apc5* is part of the platform domain which forms a base for the complex and facilitates interactions between the other domains (Sivakumar and Gorbsky 2015).

We found that CAST mice, which exhibit high penetrance of infertility in the absence of *Kif18a*, had significantly higher expression of *Apc7* in juvenile and adult testicular tissue, as well as reduced expression of *Apc5* in fetal ovarian tissue compared to B6 mice. There are coding variants within 2 Mb of *Apc5* that could impact APC5 function. In addition, there are non-coding variants that could plausibly impact *Apc5* expression, though there are presently no epigenetic profiling data to suggest that they are in or near potential gene regulatory sequences (McLaren, Gil et al. 2016). Therefore, we cannot nominate a single causative variant that would explain the observed differences in *Apc5* expression. Using publicly available structural variant call sets derived from long read *de novo* assemblies, we identified a retroviral insertion just upstream of the CAST/EiJ *Apc7* allele. Endogenous retroelements in mouse genomes are common sources of cis-regulatory variation among inbred strains, as evidenced by the 172 published phenotypes that have been attributed to endogeneous retroviral insertion alleles (Mouse Genome Database / Mouse Mine). Specifically, the endogenous RLTR10 elements are known to act as transcriptional enhancers in the male germ line and are essential regulators of spermatogenesis (Sakashita, Maezawa et al. 2020). The presence of this element could explain the increased expression of *Apc7* that we observed in CAST/EiJ males, but more data are needed to demonstrate causality. In a cell model, we found that co-depletion of KIF18A with APC7 partially relieves the mitotic arrest phenotype, which supports the transgressive allele effects that we observed in our intersubspecific F2 mouse population, where CAST/EiJ genotypes at the QTL locus are associated with higher germ cell counts, presumably due to less mitotic arrest. The concept of transgressive segregation describes traits that are observed in hybrid populations which exceed or are not observed in the parents. In laboratory mice, transgressive segregation is often observed for polygenic traits in divergent crosses. While CAST/EiJ and C57BL/6J mice are laboratory inbred strains of the same species (*Mus musculus*) - they are different sub-species (*M.m. castaneous* and *M.m. musculus*) with ∼300-500,000 years of divergence and over 20 million segregating genetic variants (Geraldes, Basset et al. 2008, Keane, Goodstadt et al. 2011, Yang, Wang et al. 2011, Phifer-Rixey, Harr et al. 2020). Since further backcrossing of the *Kif18a^gcd2^* allele onto the CAST genetic background ultimately leads to more pronounced infertility, it is likely that multiple divergent CAST alleles with additive or epistatic interactions ultimately exacerbate germ cell depletion. Quantitative trait mapping in a larger F2 population is needed to reveal the full extent of genetic variants that modulate this essential gene network in dividing cells. All nine of the candidate genes in the Chr5 QTL harbor had the common feature of harboring genetic variation that could influence gene function and exhibiting significant differential expression in at least one of the tissues / timepoints. Among these, *Kntc1*, which encodes a kinetochore protein involved in mitotic checkpoint signaling (Chan, Jablonski et al. 2000, Scaerou, Starr et al. 2001, Kops, Kim et al. 2005), stands out. While this gene was not highly expressed in fetal ovary, it was highly expressed in P5 and adult testes and it showed differential expression in B6 and CAST adult testes. Furthermore, mutations in *Kntc1* have been linked to recessive skeletal fusions, infertility, and reduced survival in mice (Fairfield, Srivastava et al. 2015). These findings underscore the potential value of further examination of *Kntc1* as genetic modifier of *Kif18a^gcd2^* phenotypes.

In animal models, the impact of APC7 depletion varies. Similar to HCT116 cells, loss of APC7 in *Drosophila* leads to mild chromosome segregation defects (Pal, Nagy et al. 2007, Wild, Budzowska et al. 2018). In humans, a loss of function founder mutation in APC7 causes the Feguson-Bonni Neurodevelopmental Syndrome (FERBON) in a large pedigree of Amish ancestry; the clinical characteristics of FERBON include neurodevelopmental delay, locomotor deficits, and premature ovarian failure (Ferguson, Urso et al. 2022). Laboratory mice engineered to carry the same *Apc7* founder mutation exhibited similar, recessive neurodevelopmental and locomotor deficits, but infertility or subfertility were not reported (Ferguson, Urso et al. 2022). These mutant mice also exhibited growth delay despite normal birth weight and adult size; a phenotype that like the recessive growth phenotype observed *Kif18A^gcd2/gcd2^* mutant mice. The Knockout Mouse Project recently reported phenotypes for a different engineered null allele of *Apc7* where, in addition to neurobehavioral, locomotor deficits and anemia, female fertility was significantly reduced in homozygous mutants (https://www.mousephenotype.org/data/genes/MGI:1929711). An engineered null allele *Apc5* (*Anapc5*^em1(IMPC)Bay^) in laboratory mice causes fully penetrant, recessive lethality prior to wean age (unpublished, https://www.mousephenotype.org/data/genes/MGI:1929722), a more severe developmental phenotype than those observed in the absence of *Kif18a* or *Apc7*. Taken together the phenotypic similarities between *Kif18a* and *Apc7* loss of function alleles further support *Apc7* as perhaps the stronger candidate gene underlying the Chr. 5 QTL, but further interrogation of these candidates in developing germ cells is needed to establish a mechanism.

In our cell models, depletion of APC7 alone did not lead to mitotic defects and the mitotic arrest phenotype of KIF18A mutant cells was partially rescued by co-depletion of APC7, in chromosomally unstable cells. Wild et al. showed that HCT116 cells depleted of APC7 exhibit slightly decreased ubiquitylation activity, but there was no significant impact on mitotic timing. Interestingly, in the same study, the phenotype of cells deficient in MAD2, part of the spindle assembly checkpoint, was partially rescued when APC7 was co-depleted. Taken together, these findings suggest that cells depleted of APC7 may be less dependent on the spindle assembly checkpoint for mitotic divisions (Wild, Budzowska et al. 2018).

Some CIN cells demonstrate phenotypes that are like those of germ cells lacking *Kif18a* function including mitotic arrest, abnormal spindle organization, and unaligned chromosomes (Czechanski, Kim et al. 2015). Sensitivity to KIF18A depletion in CIN cells is also variable depending on cell type and the contributing factors remain unknown (Marquis, Fonseca et al. 2021, Gliech, Yeow et al. 2024, Payton, Belmontes et al. 2024). Several mutations in genes encoding components of the APC have been identified in various neoplastic cell lines, and some may impair mitotic progression (Wang, Moyret-Lalle et al. 2003, Sansregret, Patterson et al. 2017, Schrock, Stromberg et al. 2020). It has been posited that these mutations allow malignant cells to slow mitosis and limit the degree of chromosomal instability, subsequently contributing to antineoplastic drug resistance (Schrock, Stromberg et al. 2020). In recent work by Gleich et al., cancer cells with lower APC activity demonstrated higher sensitivity to inhibition of KIF18A and when APC activity was increased in these cell lines, the sensitivity to KIF18A depletion was diminished. Interestingly, reduced APC activity was also implicated as a resistance mechanism to other spindle assembly checkpoint-targeting agents (Thu, Silvester et al. 2018). On review of aggregated data through DepMap, Gleich et al., identified that the top three positively correlated co-dependencies for KIF18A inhibition were APC1, APC4, and APC8. Both APC1 and 4 are part of the platform complex with APC5 and APC8 is part of the TPR lobe with APC7. These results are consistent with our findings of exaggerated mitotic arrest in cells sensitive to KIF18A depletion with co-depletion of APC5. Furthermore, consistent with our observation that co-depletion of APC7 was correlated with decreased sensitivity to KIF18A depletion in sensitive cell lines, DepMap portal data indicate a slight anti-correlation of dependency on APC7 and KIF18A across many cell lines, although this trend is not statistically significant (Tsherniak, Vazquez et al. 2017). In summary, our findings suggest that APC modulation of sensitivity to KIF18A depletion may be nuanced and can vary by APC subunit and cell type. Indeed, in studies examining *Drosophila*, differential effects are even seen with depletion of different subunits within the TPR lobe (Pal, Nagy et al. 2007).

## METHODS

### Ethics statement

All procedures involving laboratory mice were approved by The Institutional Animal Care and Use Committee of The Jackson Laboratory (under Animal Use Summary #20030). The Jackson Laboratory meets the voluntary accreditation and assessment guidelines of the American Association for Accreditation of Laboratory Animal Care International, AAALAC, a private, nonprofit organization that promotes the humane treatment of animals in science.

### Mapping cross and phenotyping

*In vitro* fertilization (IVF) was used to generate a cohort of F1 animals from CAST.129-*Kif18a^gcd/+^*cryopreserved sperm (RRID:MMRRC_034325-JAX) and oocytes from C57BL/6J (RRID:IMSR_JAX:000664) females. This work was performed by The Jackson Laboratory Reproductive Sciences Service. Genotyping for the *Kif18a^gcd2^*allele was used to identify carriers from 496 (271 males and 225 females) F1 IVF offspring. Genotyping was performed by the Jackson Laboratory Transgenic Genotyping Service using primers, methods, and reagents described on the strain data sheet (https://www.jax.org/Protocol?stockNumber=010508&protocolID=18709). Heterozygous (B6;CAST-*Kif18a^gcd2/+^*) males and females were used to set up 55 trio intercrosses, and from those 477 gonads (230 ovary, 247 testes) from B6;CAST-*Kif18a^gcd2/gcd2^* (*gcd2/gcd2*) animals and 93 (45 ovary, 48 testes) B6;CAST-*Kif18a^+/+^* (+/+) controls were collected for histological analysis at 15-18 days post-partum (dpp). Samples were fixed in Bouin’s fixative (5% saturated aqueous picric acid, 5% glacial acetic acid, 10% formalin (37-40%)) for at least 24 hours at room temperature and then transferred to the Jackson Laboratory Histopathology Services for embedding and sectioning. Briefly, samples were washed, dehydrated through ethanol, followed by xylene, and then embedded in paraffin. After facing each block, four step sections (5 uM sections, collected serially at 10 section (50 uM) intervals to avoid sampling the same cells) were placed onto glass slides. Slides were processed using a Ventana Medical Systems Discovery XT automated immunostainer. Following antigen retrieval (24 min. in Tris/EDTA at 95C), a primary rabbit polyclonal antibody against the mouse VASA homolog (MVH / DDX4) (abcam ab13840, 1:300) was used to label germ cells, followed by a secondary colorimetric detection (ChromoMap Dab, Roche # 760-159, anti-rabbit IgG). The stained slides were scanned using the NanoZoomer whole slide scanner (Hammamatsu) and FIJI / Image J image analysis software (https://imagej.net/software/fiji/) was used to quantify the area occupied by anti-MVH staining in each of 4 sections (1 image = 1 section) per F2 gonad. Since individual anti-MVH positive oocytes are discretely organized into pre-antral and antral follicles in P18 ovaries, it was possible to directly count MVH positive oocytes with the manual counting function available in ImageJ. For each animal the average area occupied by MVH staining (males) or average number of MVH positive oocytes across the 4 step sections was used to quantify the germ cell population.

### Genetic mapping

DNA samples from 225 B6;CAST-*Kif18a^gcd2/gcd2^* (*gcd2/gcd2*) F2 animals were collected for SNP genotyping. A panel of 150 SNPs that are polymorphic between C57BL/6J and CAST/EiJ and evenly spaced across all autosomes and X chromosome were selected for a genome scan, and an additional 4 SNPs (between rs3663793 [chr5:102320975 bp, GRCm38] and rs3717290 [chr5:141057331 bp, GRCm38]) were later added to narrow the initial QTL interval on Chr 5. SNP genotyping was performed by the Jackson Laboratory Genome Analysis Service (GAL). All genotype and trait data are provided in Supplementary File 1. The R/qtl quantitative trait mapping environment was used for genetic association mapping (Broman, Wu et al. 2003). Since the germ cell count data were not normally distributed, an inverse normal transformation (RankZ) was applied (Supplementary Figure 1A) and LOD scores were calculated using the Haley-Knott simple regression method with sex as a covariate (Haley and Knott 1992). Significance thresholds were determined by permutation testing (n=1,000).

### Gene expression analysis

RNA was harvested from C57BL/6J, CAST/EiJ, and B6CAST F1 fetal ovaries (11.5-13.5 dpc) and testes (5 dpp), and in each case 5 biological replicates were collected for a total of 30 samples for bulk RNA sequencing. Tissues were homogenized in TRIzol Reagent (ThermoFisher) using a Pellet Pestle Motor (Kimbal) and RNA was isolated using the miRNeasy Mini kit (Qiagen), according to manufacturers’ protocols, including the optional DNase digest step. RNA concentration and quality were assessed using the Nanodrop 2000 spectrophotometer (Thermo Scientific) and the RNA 6000 Nano Assay (Agilent Technologies). All samples had RIN values above 9. Stranded libraries were constructed using the KAPA mRNA HyperPrep Kit (Roche Sequencing and Life Science), according to the manufacturer’s protocol. Briefly, the protocol entails isolation of polyA containing mRNA using oligo-dT magnetic beads, RNA fragmentation, first and second strand cDNA synthesis, ligation of Illumina-specific adapters containing a unique barcode sequence for each library, and PCR amplification. The quality and concentration of the libraries were assessed using the D5000 ScreenTape (Agilent Technologies) and KAPA Library Quantification Kit (Roche Sequencing and Life Science), respectively, according to the manufacturers’ instructions. Libraries were sequenced 100 bp paired-end on an Illumina HiSeq 4000 using HiSeq 3000/4000 SBS Kit reagents. Additional RNASeq data from C57BL/6J and CAST/EiJ adult (16 weeks) testes were obtained from GEO (GSE235498: GSM7503634, GSM7503643, GSM7503649, GSM7503650, GSM7503636, GSM7503646, GSM7503653).

FastQC (version 0.11.33) was used to assess the demultiplexed reads prior to alignment. Since the CAST/EiJ, *Mus musculus castaneous* sub-species genome is significantly diverged from the mouse reference genome (C57BL/6J, *Mus musculus domesticus*), reads were aligned using EMASE, Expectation-Maximization (EM) algorithm for Allele Specific Expression (ASE). This algorithm accounts for strain-specific genetic variation by aligning to transcriptomes adjusted for known genetic variation, and estimates transcript abundances by allele and by gene (Raghupathy, Choi et al. 2018). Estimated read counts per sample across all samples were used as input for differential expression analysis using R/Deseq2 (Love, Huber et al. 2014). Sample metadata for all RNAseq samples are in Supplementary Table 1. Individual sample ‘read count per transcript’ (**_pe_ec_emase-zero_m2.counts) files were loaded into R and count matrices were generated by creating a new data frame with transcript IDs as rows and sample counts per sample as columns. Two count matrices, one for ovary samples and one for testis samples were used for downstream analyses. PCA (prcomp in baseR) was computed across all samples to detect outliers (beyond 2 standard deviations of PC1 or PC2) and none were found. Ggplot2 was used to generate plots of read counts per transcript, gene, and sample. For differential expression analysis, DESeqDataSecFromMatrix was used with each count matrix providing counts, coldata to sample_metadata (Supplemental Table 1), and design ∼ strain. Expressed genes (baseMean>10) with adjusted with adjusted p-value (padj) of less than 0.05 were scored as differentially expressed.

### Candidate gene analysis

The approximate Bayesian interval for the Chr5 QTL was defined using the command *bayesint* in R/QTL (Broman, Wu et al. 2003) with a probability threshold of 70%. This interval was defined by top scoring SNP (rs3670250, lod score = 6.09, chr5:119,982,112 (GRCm38)) and the flanking markers (rs3719351 at chr5:113,989,413 and rs3664741 at chr5:125,025,240 (GRCm38)). All genes with human orthologs in this QTL interval were filtered to only those expressed (mean estimated read count greater than 10) in fetal ovaries (12.5-13.5 dpc) and testes (5 dpp), as well as adult testis. Using published whole genome variant data for CAST/EiJ from the Mouse Genomes Project (Lilue, Doran et al. 2018) (obtained from GenomeMUSter (Ball, Bogue et al. 2023)), the gene list was annotated for the presence of protein damaging (missense, stop gain, stop lost, splice acceptor / donor site variants (SNPEff annotation of ‘Medium’ or ‘High’). Using the Mouse Genome Database (Blake, Baldarelli et al. 2021) and MouseMine (Motenko, Neuhauser et al. 2015), and the associated phenotype (Mammalian Phenotype Ontology) and GOSlim functional annotations were assembled for the remaining genes. The candidate gene list was further annotated for GOSlim and Mammalian Phenotype Ontology terms related to *Kif18a* functions and phenotypes as follows: GOSlim terms: GO:0046785 microtubule polymerization, GO:0042127 regulation of cell population proliferation, GO:0045840 positive regulation of mitotic nuclear division, GO:0007094 mitotic spindle assembly checkpoint signaling, GO:0005828 kinetochore microtubule, GO:0000776 kinetochore, GO:0000775 chromosome, centromeric region, GO:0005694 chromosome, GO:0000070 mitotic sister chromatid segregation, GO:0000278 mitotic cell cycle, GO:0000086 G2/M transition of mitotic cell cycle, GO:0051301 cell division, GO:0007049 cell cycle; Mammalian Phenotype Ontology: MP:0011100 preweaning lethality, complete penetrance, MP:0001926 female infertility, MP:0011110 preweaning lethality, incomplete penetrance, MP:0002685 abnormal spermatogonia proliferation, MP:0004901 decreased male germ cell number, MP:0011094 embryonic lethality before implantation, complete penetrance, MP:0011098 embryonic lethality during organogenesis, complete penetrance, MP:0001119 abnormal female reproductive system morphology, MP:0001922 reduced male fertility, MP:0001923 reduced female fertility, MP:0011092 embryonic lethality, complete penetrance, MP:0011101 prenatal lethality, incomplete penetrance, MP:0011102 embryonic lethality, incomplete penetrance, MP:0011108 embryonic lethality during organogenesis, incomplete penetrance, MP:0013292 embryonic lethality prior to organogenesis.

### Cell Culture and knockdown

OVCAR3 and RPE1 cells were purchased from ATCC. HeLa Kyoto cells were a generous gift from Ryoma Ohi’s lab at the University of Michigan and SKOV3 cells were a generous gift from Alan Howe’s lab at the University of Vermont. HeLa Kyoto and RPE1 cells were cultured in Minimum Essential Medium Eagle Alpha (Gibco) supplemented with 10% Fetal Bovine Serum (Gibco). SKOV3 cells were cultured in McCoy’s 5A Modified Media (Gibco) supplemented with 10% Fetal Bovine Serum (Gibco). OVCAR3 cells were cultured in RPMI 1640 medium (ATCC) supplemented with 20% Fetal Bovine Serum (Gibco), 0.01mg/mL Bovine insulin (Sigma-Aldrich), and 10% Penicillin-streptomycin (Gibco). All cell types were incubated at 37C with 5% CO2.

To inhibit proteins of interest, cells were transfected with siRNA using Lipofectamine RNAiMAX Transfection Reagent (Invitrogen) in Opti-MEM Reduced-Serum Media (Gibco). HeLa Kyoto and RPE1 cells were treated with 5 pmol siRNA. SKOV3 and OVCAR3 cells were treated with 10pmol ANAPC5 and ANAPC7 siRNA, as a higher concentration was required to achieve adequate knockdown. Specific siRNAs used include pools of Silencer and Silencer Select KIF18A (Invitrogen; GCUGGAUUUCAUAAAGUGGtt, CGUUAACUGCAGACGUAAAtt), SMARTPool ANAPC7 (Dharmacon; GCAAUGGACCAGUAUAGUA, UCGCUUAGAUUGUUAUGAA, GUUCAAGCUCUGCUACUUA, GGAAUGCUGUGAGUAAGUA), pools of Silencer Select ANAPC5 (Life Tech s28130, s28131, and s28129; GAAGGCGAGUUGAAGGAUATT, GGAAAGCGGUUGUAUUACATT, GCACUUGAAGGAACGAUUUTT), ON-TARGETplus non-targeting pool (Dharmacon; UGGUUUACAUGUCGACUAA, UGGUUUACAUGUUGUGUGA, UGGUUUACAUGUUUUCUGA, UGGUUUACAUGUUUUCCUA), and pools of Silencer and Silencer Select negative controls (Invitrogen Catalog #4390843 and #AM4613). For double knockdowns involving the inhibition of two proteins, Lipofectamine RNAiMAX was used at a lowered concentration (0.7× the concentration used for single knockdowns) to mitigate toxicity.

### Immunofluorescence

Cells were grown on glass coverslips and were fixed using 4% paraformaldehyde in −20 °C methanol. Cells were then blocked with 20% goat serum in antibody diluting buffer (Abdil-TBS, 1% BSA, 0.1% Triton X-100, and 0.1% sodium azide). Cells were then incubated with the following primary antibodies in Abdil; mouse anti-α-tubulin (DM1α) 1:1000 (Millipore Sigma, Cat no. T6199), Rabbit anti-KIF18A 1:100 (Bethyl Laboratories, Cat no. A301-080A), and Guinea pig anti-CENP-C 1:250 (MBL) for one hour at room temperature. Cells were then incubated with secondary antibodies conjugated to Alexa Fluor 488, 594, and 647 at concentrations of 1:500 for one hour at room temperature. Coverslips were then mounted onto glass slides using Prolong Gold anti-fade mounting medium with DAPI.

### Microscopy

Fixed-cell images were acquired using a Ti-E or Ti-2E inverted microscope (Nikon Instruments) driven by NIS Elements software (Nikon Instruments). Images were captured using a Clara cooled charge-coupled device (CCD) camera (Andor). Fixed-cell images were taken using a Plan Apo 40X, 0.95 NA objective (Nikon).

### Western Blot

Cells were lysed in PHEM lysis buffer (60 mM Pipes, 10 mM EGTA, 4 mM MgCl2, and 25 mM HEPES) with 1% Triton X-100 and protease inhibitors, incubated on ice for 10 minutes, and then centrifuged at maximum speed for 10 minutes. Laemmli buffer with β-mercaptoethanol was added to the supernatant and samples were boiled at 95 °C for 10 minutes. Lysates were run on 4–15% gradient gels (BioRad), transferred to a PVDF membrane (BioRad), and blocked for 1 hour in 1:1 Odyssey Blocking Buffer (Li-Cor) and TBS. Membranes were incubated with primary antibodies overnight at 4 °C. Primary antibodies included 1:1000 mouse anti-GAPDH (Millipore, Cat no. MAB374), 1:500 rabbit anti-KIF18A (Bethyl Laboratories, Cat no. A301-080A), 1:2000 rabbit anti-ANAPC7 (Bethyl Laboratories Cat no. A302-551A), and 1:1000 rabbit anti-ANAPC5 (Bethyl Laboratories Cat no. A301-026A.). Secondary antibodies included goat anti-rabbit IgG DyLight 800 conjugate (Invitrogen, Cat no. SA5-10036) and goat anti-mouse IgG DyLight 680 (Invitrogen, Cat no. SA5-10170), diluted to 1:10,000 in 1:1 Odyssey blocking buffer/TBS and added to the membrane for one hour at room temperature. Membranes were imaged using an Odyssey CLx (Li-Cor).

### Mitotic timing and mitotic index analyses

Mitotic index was measured using fixed-cell images by counting the number of mitotic cells divided by the total number of cells. All mitotic index fields were taken with a 40x objective. Contingency tables were created in Prism 10 to compare the total number of cells that either divided or failed to divide; statistical significance was then determined using Chi-square tests to compare between the conditions within each cell line.

### Knockdown quantification analysis

The efficiency of siRNA-mediated protein knockdowns was measured via quantitative densitometry analyses of western blots in ImageJ.

### Statistics and reproducibility

Data are represented as mean ± standard deviation (SD), unless otherwise specified. All statistical tests for RNAi experiments were performed using GraphPad Prism version 10.1.0. Statistical test for genetic mapping and RNASeq were performed using R/QTL and R/Deseq2.

## ACKNOWLEDGEMENTS

We gratefully acknowledge the contribution of the Genome Analysis, Reproductive Sciences, and Histopatholgy Services at The Jackson Laboratory for their expert assistance with the SNP mapping panel design, *in vitro* fertilization, and automating immunostaining respectively. The Scientific Services at the Jackson Laboratory are partially supported by the National Cancer Institute under award number P30CA034196. Additionally, the research reported in this publication was supported by a grant from the National Institutes of Childhood Health and Disease (R03HD078485 to L.G.R), a pilot award from the Cancer Center at The Jackson Laboratory (P30CA034196 to L.G.R. and J.S.), and an Outstanding Investigator Award (R35GM144133 to J.S.). The content is solely the responsibility of the authors and does not necessarily represent the official views of the NIH.

## AUTHOR CONTRIBUTIONS

Investigation: CN, WM, AC, CB. Formal analysis: CN, WM, NR, AF, JS, LGR. Visualization: CN, WM, NR, AF. Methodology: JS, LGR. Conceptualization: JS, LGR. Project administration: JS, LGR. Writing – original draft: CN, LGR. Writing – review & editing: JS, LGR. Supervision: JS, LGR. Funding acquisition: JS, LGR

## DATA AVAILABILITY

Illumina RNASeq data are available through GEO (GSE281149).

## ADDITIONAL INFORMATION

The authors have no competing interests

---

**Supplementary Figure 1.**
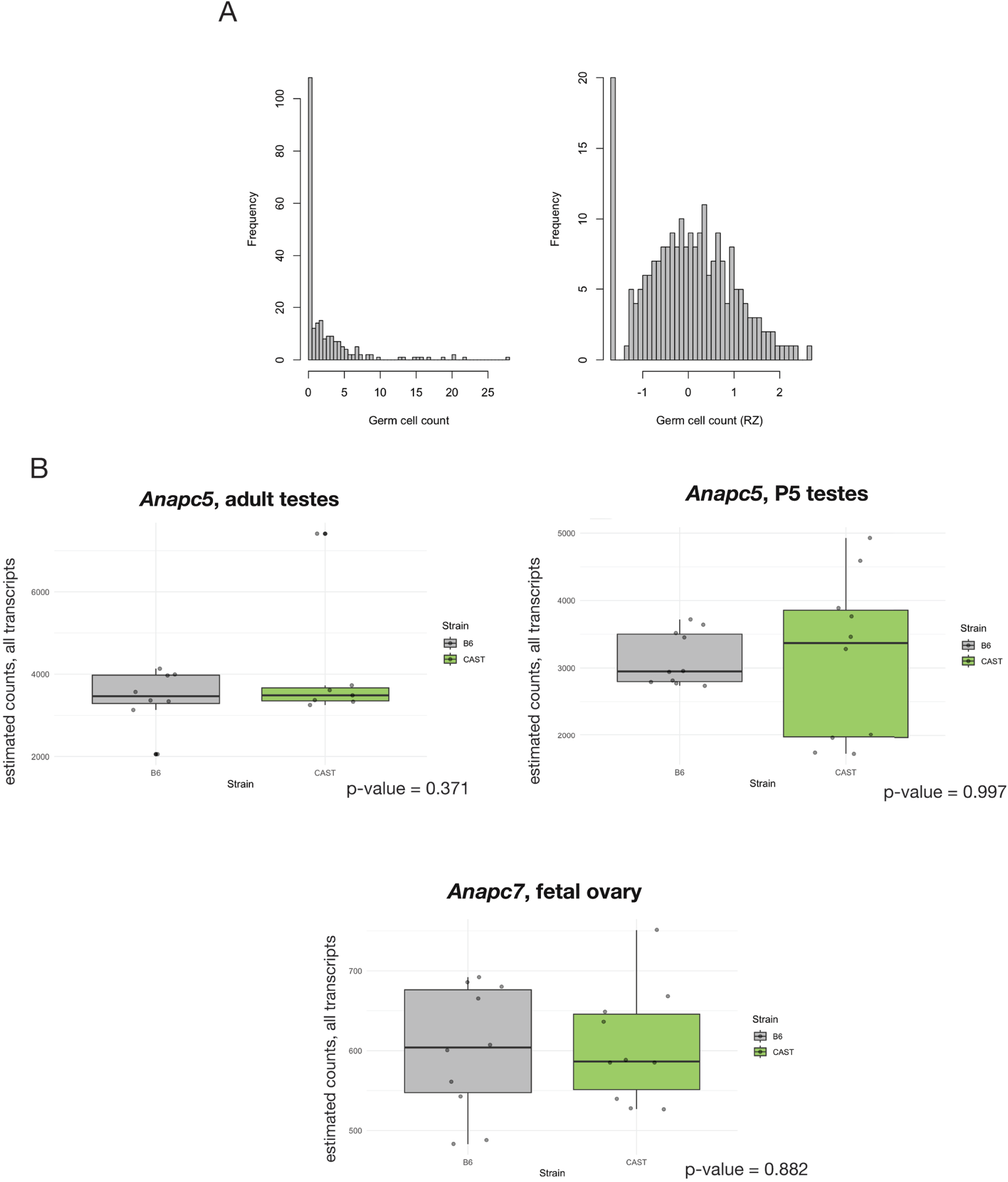
Variation of mouse germ cell populations in B6;CAST-*Kif18a^gcd2/gcd2^* F2, raw data and RankZ transformed (A). Tissues and timepoints where *Anapc5* and *Anapc7* expression differences between B6 and CAST were not significant, n=5 for each tissue type/strain (B).

**Supplementary Figure 2.**
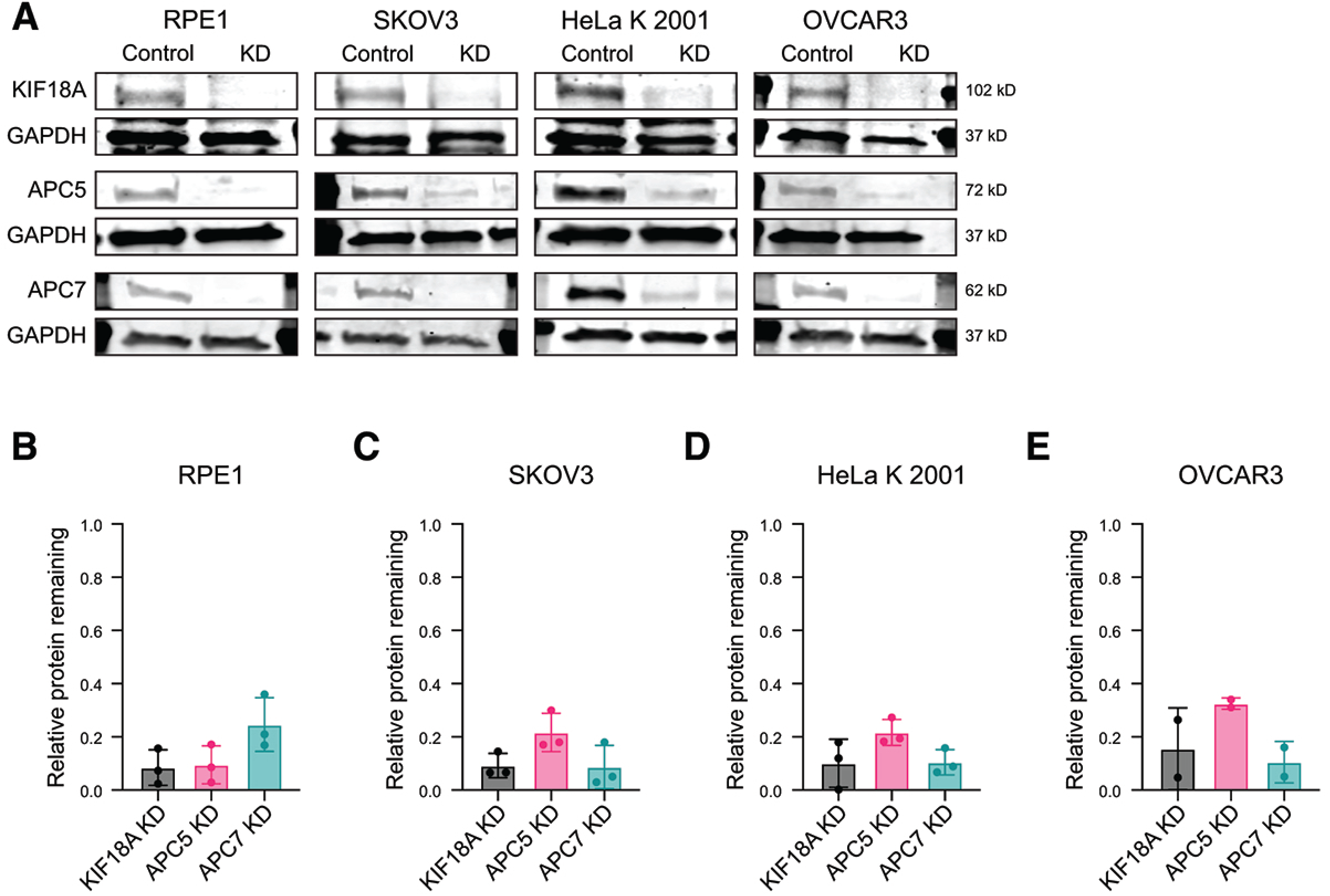
Representative Western blots of proteins of interest following knockdown (KD) with siRNA in RPE1, SKOV3, HeLa, and OVCAR3 cells (3 independent experiments for each protein for RPE1, SKOV3, and HeLa cells, 2 independent experiments for each protein for OVCAR3 cells.) (A) B,C,D,E Quantification of siRNA KD in RPE1 (B), SKOV3 (C), HeLa (D), and OVCAR3 (E) cells measured via Western blot and normalized to control siRNA condition and GAPDH loading control. n=3 individual 3 independent experiments for each protein for RPE1, SKOV3, and HeLa cells (all conditions), 2 independent experiments for each protein for OVCAR3 cells (all conditions).

## SUPPLEMENTARY FILES AND TABLES

**Supplementary File 1.** Sample genotype and germ cell trait data from F2 progeny

**Supplementary Table 1.** Metadata for RNASeq samples and files

**Supplementary Table 2.** Candidate gene analysis based on expression, B6 vs. CAST variation, and gene annotations (function and phenotype).

**Supplementary Table 3.** Differential gene expression analysis of all candidate genes within the Chr 5 QTL across three tissues / timepoints.

